# Mechanical and morphological features of the cockroach antenna confer flexibility, reveal a kinematic chain system and predict strain information for proprioception

**DOI:** 10.1101/2025.04.07.647640

**Authors:** Lingsheng Meng, Parker McDonnell, Kaushik Jayaram, Jean-Michel Mongeau

## Abstract

A broad class of animals rely on touch sensation for perception. Among insects, the American cockroach *P. americana* is a touch specialist that uses a pair of soft antennae with distributed sensors to touch its environment to guide decision making. During touch, forces on the antenna can activate thousands of mechanosensors. To understand the content of this sensory information, it is critical to understand how antenna mechanics shape the transmission of contact forces. Here, we investigate the mechanical behavior and morphology of the American cockroach antenna at the individual segment level through experiments, mathematical modeling, imaging with Micro-Computed Tomography (Micro-CT), 3D reconstruction of antenna morphology, and finite element modeling (FEM). Our experimental results and model predictions reveal that the antenna flagellum bends according to a kinematic chain model, with rigid segments connected by joints. Whereas the middle region of the antenna consistently fractured under cyclic bending, the tip region remained intact under large deformations, revealing mechanical specialization along the antenna. Micro-CT imaging revealed an invagination of the exocuticle at segment intersections of the tip. To test the hypothesis that this structure can enhance flexibility and robustness, we used FEM and confirmed that the invagination allows for larger bending without structural failure (buckling). Applying FEM to a morphologically accurate kinematic chain model of the flagellum revealed the relationship between the local strain at the location of marginal sensilla and intersegment angle, predicting the information available for antenna proprioception. Taken together, these findings reveal biomechanical adaptations of insect antennae and provide a critical step toward a mechanistic understanding of touch sensation in a touch specialist.

**SUMMARY STATEMENT:** By combining experiments, imaging and modeling, we demonstrate distinct mechanical features in the cockroach antenna and provide a framework to model the neuromechanics of the antenna in an insect touch specialist.

## INTRODUCTION

All insects have antennae, except for a primitive hexapod (Protura) (Schneider, 1964). Antennae serve critical sensory functions, providing sensory feedback across modalities, including touch, gustation, olfaction, magnetoception, etc. The remarkable diversity of insect antennae can enable the discovery of unique biological adaptations and inspiration for the design of new sensors. Among insect antennae, the filiform (thread-like) antennae of the American cockroach *Periplaneta americana*—a touch specialist—plays a vital role in sensory perception for guiding locomotion. Cockroaches such as *P. americana* rely on their antennae for distinct touch-mediated behaviors, including tactile exploration and rapid evasive maneuvers, such as wall following (Camhi and Johnson, 1999; Comer et al., 2003; Cowan et al., 2006; Okada and Toh, 2000; Okada and Toh, 2006). To function effectively, the antenna must remain compliant during contact while maintaining structural integrity during non-contact episodes (Rajabi et al., 2018). For instance, the tapered profile of the antenna enhances its ability to bend and comply to mechanical stimuli at the tip (Mongeau et al., 2014). The resulting decreasing flexural stiffness along its length increases preview distance, which is particularly critical for feedback control during high-speed running (Cowan et al., 2006). The passive mechanics and structure of the American cockroach flagellum appears important for sensing and locomotion control: it rapidly damps perturbations (within one stride cycle), maintains contact after impact, and the embedded hair sensilla play a mechanical role in reconfiguring the antenna to enhance control (Mongeau et al., 2013; Mongeau et al., 2014). Further, neural processing within the antenna appears to be well suited to implement feedback control during rapid running maneuvers (Lee et al., 2008; Mongeau et al., 2015). However, it remains unclear how local structural properties of the flagellum shape mechanosensory information for tactile sensing and perception. For instance, marginal sensilla embedded within flagellar segments are thought to act like strain gauges, yet it is unknown how strain develops locally within segments during natural bending. Estimating the local cuticular strain—a prerequisite to understand the neuromechanics of touch—requires accurate models of antenna mechanics.

While the filiform antenna of crickets and cockroaches have been approximated as a continuous tapered beam, these antennae are segmented, which could have important implications for how strain and stresses develop during natural bending. Mechanical measurements and modeling of insect antennae have revealed important insights into their structural properties. For instance, detailed Finite Element Models (FEM) have been developed for the stick insect *Carausius morosus*. This model incorporated a tapering radius, variations in cuticle thickness, the presence of compliant endocuticle, and the segmented annuli or flagellomeres of the antenna (Rajabi et al., 2018) (henceforth referred as “segment”). The tapered shape of the stick insect antenna seems to have the most significant impact on static bending, as also described in cockroaches (Mongeau et al., 2014), while the compliant endocuticle primarily influences the dynamic response (damping) (Rajabi et al., 2018). With approximately 150 segments composing the American cockroach antenna (Schafer and Sanchez, 1973; Schafer and Sanchez, 1976), the role of individual segments and their intersections in influencing bending responses remain poorly understood. Interestingly, tensile testing experiments in the American cockroach showed that the antenna breaks at segment intersections (Donley et al., 2022), suggesting that these features could be an area of high stress concentration during transverse bending. Further, in the cricket *Acheta domesticus* the antenna appears to shorten while bending (Loudon et al., 2014), suggesting that soft cuticular joints are important for antenna articulation. To close the gap in our understanding of how these structural features contribute to the antenna’s overall mechanical performance, here we explore the mechanical behavior of the antennae at the segment level using a combination of animal experiments, analytical modeling, simulation, and imaging combined with 3D morphological reconstruction.

To investigate the antennal segment response during large, naturalistic deformation, we measured the bending response of the flagellum experimentally. We investigated the morphology of flagellar segments across the base, middle, and tip regions using Micro-Computed Tomography (micro-CT). By comparing an analytical model to experimental data, we determined that the antenna flagellum can be accurately modeled as a kinematic chain. By combining imaging and FEM, we discovered an invagination between segments in the tip region that enhances compliance during large bending. By generating morphologically accurate models of the segments from micro-CT, we estimated the strain field near the location of marginal sensilla, which are important sensors for joint proprioception. Taken together, this work provides a critical step toward developing a detailed neuromechanical model of an insect antenna, which could yield a deeper mechanistic understanding of touch sensation.

## MATERIALS & METHODS

### Antenna cyclical bending experiment

To investigate the mechanical response of flagellar segments under bending, we tethered a male cockroach onto a custom 3D printed holder. A custom fixture was designed to fix the antenna at specific regions along its length, enabling recordings from the base, middle, and tip (**Figure 1A**). For cyclical bending experiments, we defined the base as segments ∼10-50, the middle as segments ∼70-100, and the tip as segments ∼130-150, which was based on morphological distinctions along the length, e.g. the rate of change of segment length from base to tip (Kapitskii, 1984; Mongeau et al., 2014). We cyclically bent the antenna manually, approximately three segments away from the fixed region (**Movie 1**). The bending limit was determined by the shape of the custom fixture (with 0.4 mm curvature) and the mechanical failure point of the antenna. The entire cyclical bending process was captured using a high-resolution microscopy camera (CS165CU-Zelux 1.6 MP Color CMOS Camera) mounted on a stereomicroscope. To analyze the maximum bending angles between segments during cyclical bending, we developed custom MATLAB (MathWorks, Inc., Natick, Massachusetts, United States) code for manually labeling segments. In frames where the antenna reached its maximum bending position (**Figure 1A**), we manually labeled the proximal and distal midpoint of two adjacent segments. Using these labeled points, we calculated the angle between two segment vectors. For each trial, the average maximum bending angle across all frames was computed.

**Figure 1.**
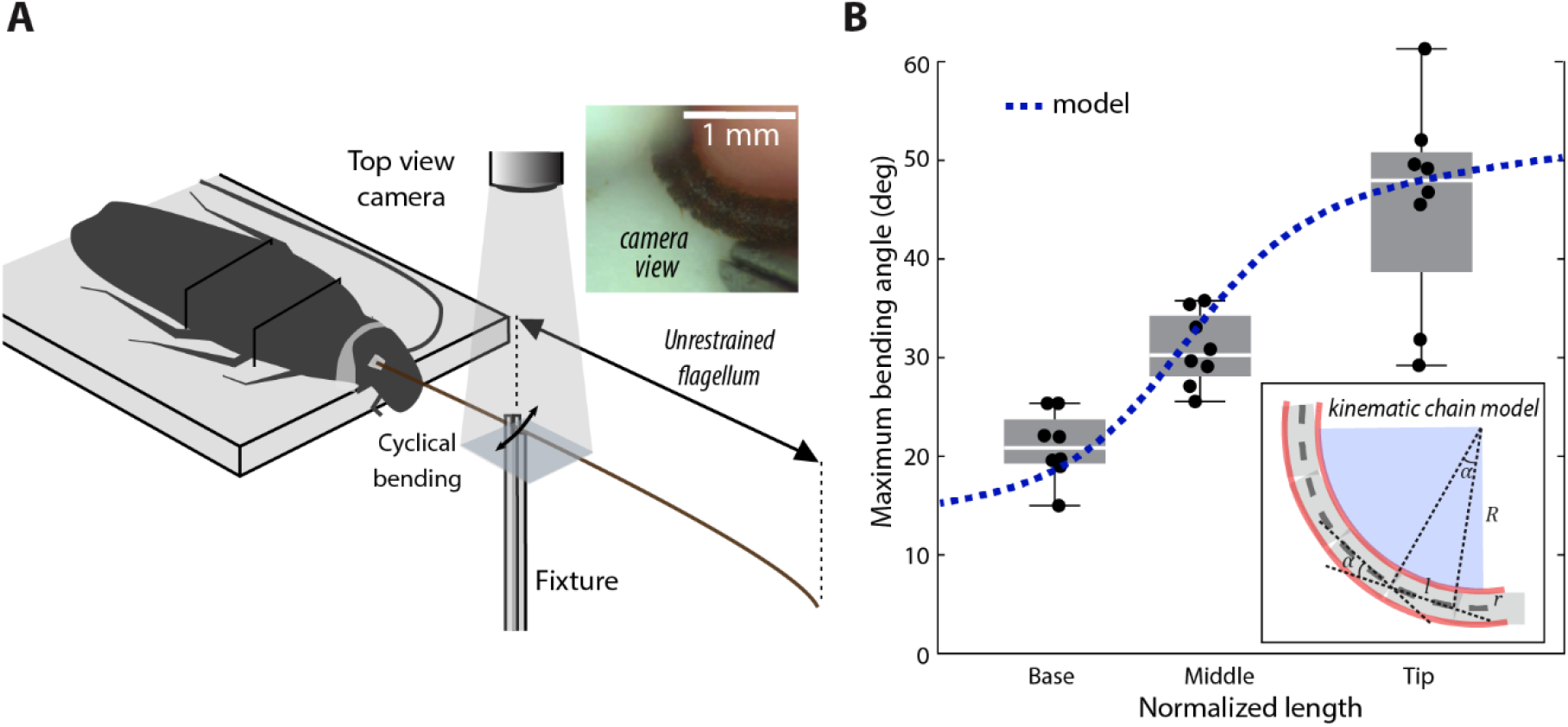
Cyclical antenna bending experiment and kinematic chain model. (A) Experimental setup. Inset: segments during bending in camera view. (B) Box plot: empirical maximum segment angles during bending. Blue dashed line: analytical prediction of bending angles from a kinematic chain model. Right bottom: analytical model, where *l* is the segment length, *R* is the curvature radius of the surface, *r* is the segment radius, and *α*is the bending angle between adjacent segments.

### Analytical calculation of maximum bending angles between segments

To estimate the maximum bending angles between segments, we derived an analytical expression for the local curvature of a kinematic chain (**Figure 1B**). The effective radius of curvature was defined as the sum of the surface curvature radius and the segment radius. We further assumed that the angle between segment was evenly distributed along the antenna length at a local scale (see Supplement). Under these assumptions, the bending angle α between adjacent segments was estimated using the following relationship (see Supplement for derivation):

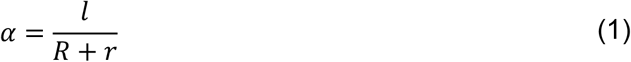

where *α* is the angle between adjacent segments, *l* is the segment length, *R* is the curvature radius of the surface (0.44 mm), and *r* is the segment radius. The segment length and radius along the length of the antenna were measured in previous work (Mongeau et al., 2014). This analytical approach provides a theoretical baseline for comparing the experimentally observed segment angles during bending.

### Micro-CT scanning of antenna regions

To examine the morphology of antenna segments in the base, middle, and tip regions, we performed micro-CT scanning. After sedating a male cockroach by chilling it for approximately 30 minutes, we cut the antenna at the head-scape joint. We then prepared the antenna based on the following dehydration and fixation protocol (Alba-Tercedor et al., 2021). First, the freshly cut antenna was immersed in distilled water for 1 hour for cleaning. Dehydration followed, using a graded ethanol series: 50%, 70%, 90%, and 3 changes of 100% ethanol, with each step lasting 20 minutes. After dehydration, the middle and tip samples were fixed in a 1% iodine solution in 100% ethanol for 48 hours to enhance the visibility of soft tissues during micro-CT imaging. Following fixation, the sample was immersed in hexamethyldisilazane (HMDS) for 2 hours. The sample was then air-dried for at least 12 hours before scanning. The sample was divided into 3 sections representing the base, middle, and tip regions. Each section was trimmed and glued to a custom-designed holder. Micro-CT scans of these 3 sections were performed using a Zeiss Versa 620 with a spatial resolution of approximately 2 μm. For microCT, the base region comprised the 11^th^ to 13^th^ segments, the middle region included the 81^st^ to 83^rd^ segments, and the tip region consisted of the 141^st^ to 143^rd^ segments, counting from the scape (1^st^ segment of the antenna).

### Finite element modeling of antenna segments

We developed three region-specific finite element models representing the base, middle, and tip regions of the antenna. We utilized a colormap in Dragonfly software (Comet Technologies Canada Inc., Montreal, Canada) to distinguish the exocuticle and endocuticle from micro-CT imaging data. Cross-sectional images of the antenna, perpendicular to its long axis, were exported for segmentation of the cuticle. These cross-sections were then annotated using a custom MATLAB script, where two ellipses were drawn to delineate the boundaries of the exocuticle (blue ellipse in **Figure 2B**) and endocuticle (red ellipse in **Figure 2B**) layers. The parameters of the ellipse, including the rotation center and lengths of the long and short axes, were used to quantify the variation in the radii of the exocuticle and endocuticle across the scanned segments. The thickness of the exocuticle was determined by calculating the difference between the axes of the inner and outer ellipses. As the thickness showed little variation within each segment (**Figure 2B, Fig. S2**), a constant thickness was chosen based on the average value: 6 μm for the base region, and 4 μm for both the middle and tip regions. Since the sgifness of the outer exocuticle is significantly higher than the endocuticle, we expect it to play a dominant role in the static mechanical deflection of the antenna (Rajabi et al., 2018), and selected it for constructing finite element models.

**Figure 2.**
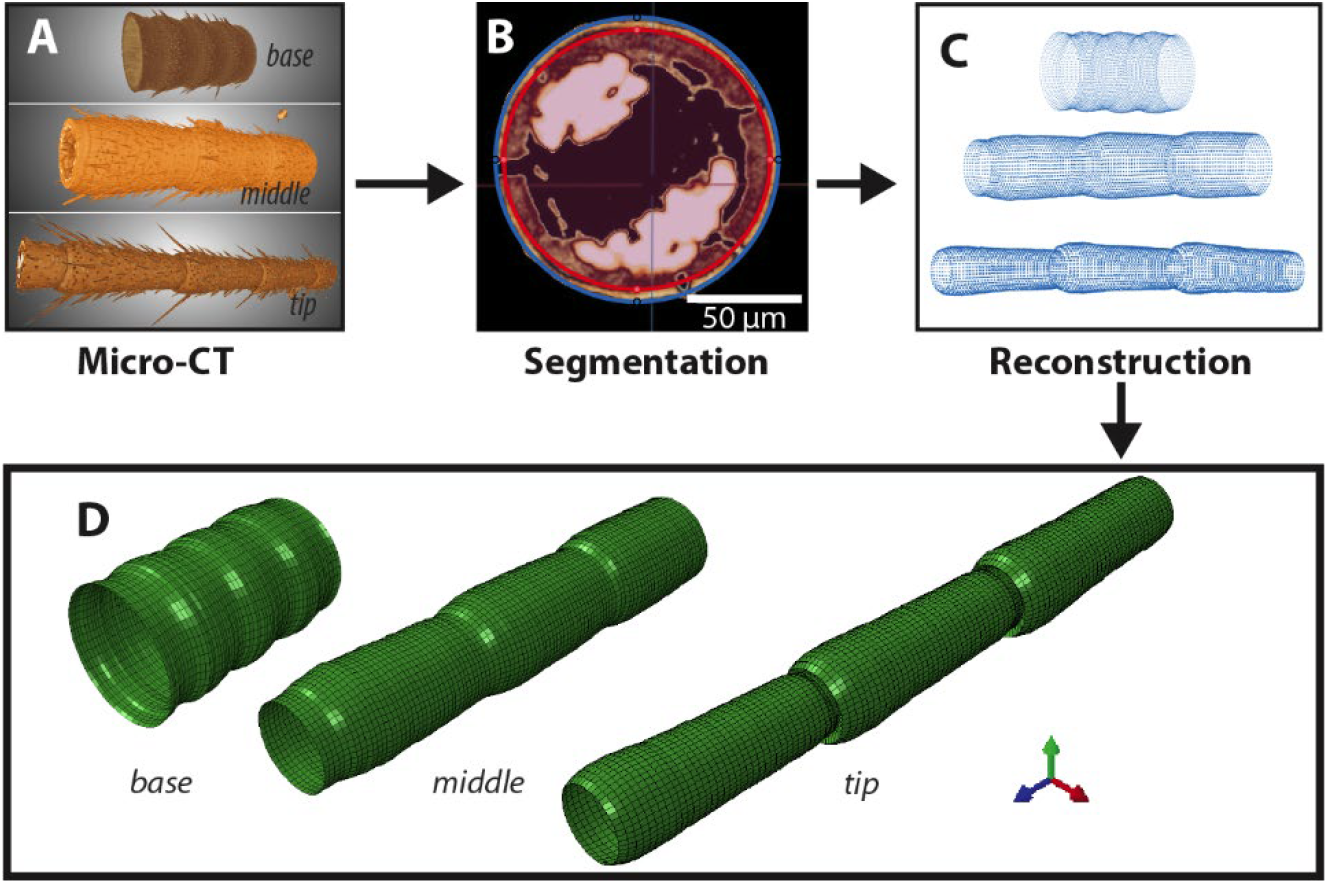
Methods to generate finite element models based on micro-CT imaging. (A) Micro-CT imaging of base, middle and tip regions of the antenna. (B) Example segmentation of cross-section of the tip region of the antenna. Blue ellipse: segmented outline of the exocuticle. Red ellipse: segmented outline of the endocuticle. (C) The point cloud generated from segmentation used to reconstruct the surface of the antenna. (D) Finite element models were constructed by morphing the coordinates of nodes to match the point cloud data.

General purpose, four-node shell elements with reduced integration (S4R) were employed to generate the mesh, as the thickness of the exocuticle is approximately two orders of magnitude smaller than the radius of the exocuticle. Point clouds were generated from the segmentations, with each point corresponding to a node in the finite element model (**Figure 2C**). The initial baseline finite element model, modeled as a thinned-walled cylinder using S4R elements, was created in Abaqus (Dassault Systèmes, Rhode Island, United States) with the same number of nodes as the points in the corresponding point cloud. The coordinates of the nodes were then morphed to match the coordinates of the point cloud (**Figure 2D**). Since we observed unique invagination at the intersections of segments in the tip region, the mesh density in the tip region’s finite element model was refined to accurately capture this feature. The detailed parameters and material properties for the finite element models of three antenna regions—base, middle, and tip—are summarized in Table 1 and based on previous estimates of insect cuticle (Combes and Daniel, 2003; Rajabi et al., 2018; Vincent and Wegst, 2004; Wainwright, 2020).

**Table 1.** Parameters of the finite element models.

| | Elem. No. | Node No. | Thickness<br>($\mu\text{m}$) | Modulus<br>(GPa) | Density<br>($\text{kg/m}^3$) | Poisson's<br>Ratio |
| --- | --- | --- | --- | --- | --- | --- |
| Base | 3200 | 3250 | 6 |  |  |  |
| Middle | 4950 | 5000 | 4 | 9.6 | 1200 | 0.49 |
| Tip | 5950 | 6000 | 4 |  |  |  |

### Pressure and hemolymph influence during bending

In our finite element model, the antenna was treated as a hollow tube. However, the antenna contains hemolymph, trachea, and other internal structures that may influence its mechanical behavior. Hemolymphatic pressure has been shown to provide structural support in the cockroach antenna during tensile testing to failure (Donley et al., 2022). The cockroach antenna contains hemolymphatic vessels with openings that allow hemolymph to flow through the antenna cavity (hemocoel) (Kapitskii, 1984). Interestingly, in our bending experiments on freshly cut antennae, no visible hemolymph leakage was observed from the cut cross-section. If hemolymphatic pressure plays an important role in antenna bending, we would expect a release of pressure and substantial hemolymph leakage during bending. The absence of such leakage suggests that hemolymphatic pressure may have a minimal effect on segmental bending behavior. Considering the effect of interior pressure on a hollow tube, both the maximum bending moment and the critical curvature of the tube increase by the same factor 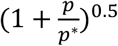, where *p* is the internal pressure, and *p*^*^is the characteristic pressure determined by the geometry and material properties of the tube (Calladine, 1983). More specifically,

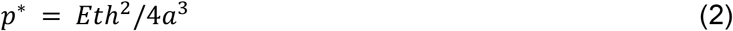

where *E* is the elastic modulus, *t* is the shell thickness, *h* is the effective shell thickness, and *a* is the radius of shell. Using the geometry and material properties of the base region, we estimated *p*^*^ to be approximately 85 kPa. Thus, if *p* is small relative to *p*^*^, we would expect a negligible contribution of hemolymph pressure. During resting, the hemolymphatic pressure in the hemocoel of insects appear to be on the order of ∼1 kPa or less, thus we would expect a negligible contribution, notwithstanding the possible role of the antenna accessory pulsatile organs in modulating pressure (Pass, 2000). Therefore, we did not include the influence of hemolymphatic pressure in our model.

### Statistics

We report mean ± 1 standard deviation, unless otherwise specified. When displaying box plots, the central line represents the median, the box edges denote the 25th and 75th percentiles, and the whiskers extend to the most extreme data points.

## RESULTS

### Flagellar segment interactions during bending are predicted by a kinematic chain model

We hypothesized that segment interactions during large bending could be predicted by a kinematic chain model, which assumes that the antenna is a series of rigid links (segment) connected by flexural joints (Demir et al., 2010). To test this hypothesis, we mechanically deflected the antenna to reproduce naturalistic large bending, such as during grooming. During grooming cockroaches uses their front leg to fold the antenna at the base with high curvature (Böröczky et al., 2013). We measured segmental movement under large cyclical bending with constant curvature. The antenna was cyclically bent to conform to a surface with a constant radius of curvature of 0.4 mm, which is approximately half the estimated minimum radius of curvature radius during grooming (see Supplement).

In all regions (base, middle, and tip), the segments folded inward during bending (**Movie 1, 2, 3**). The degree of folding and segment responses varied across regions. Specifically, the individual limit of bending angles increased progressively from the base to the tip (**Figure 1B**). The average maximum bending angles were statistically different among the base, middle, and tip regions (ANOVA, *p* = 1.5e-6, *n* = 8 antennae, *N* = 4 animals). Consistent with experimental data, analytical predictions of bending angles based on a kinematic chain model predicted an increasing trend from the base to the tip (blue dashed line in **Figure 1B**).

Distinct regions of the antenna had distinct responses to large bending. In the base region, the shorter segment length and larger radius resulted in smaller bending angles for each joint. Additionally, due to the higher number of segments over the same bending length, there was greater freedom of movement, leading to reduced bending angles for individual joints. Notably, the middle region was prone to breakage during the experiments before reaching the given curvature (**Movie 2**), leading to maximum bending angles smaller than the analytical predictions. In contrast, the tip region consistently withstood larger bending without structural failure. Agreement between data and a simple analytical model suggests that, for the same curvature, the bending angle of each segment may be governed primarily by the segment’s length and radius. Interestingly, the kinematic chain model did not fully predict the ultimate fracture bending angle, suggesting more complex interactions between segments, particularly towards the tip.

### Micro-CT scans reveal regional specialization across the flagellum

During the antenna bending experiments, we observed the segments folded inward in all three regions. The tip region demonstrated a remarkable capacity to withstand large bending without failure. To gain deeper insight into the 3D morphology of the segment intersections, we conducted micro-CT scans of the base, middle, and tip regions. We obtained morphological data, including radius, segment length, and thickness, through manual segmentation of Micro-CT data (**Figure 3A, B**). Consistent with previous morphological measurements (Kapitskii, 1984; Mongeau et al., 2014), the base region had a significantly larger average radius (199.09 ± 6.58 μm) compared to the middle (106.10 ± 4.90 μm) and tip (63.64 ± 5.03 μm) regions. Conversely, the segment lengths in the base region (194.05 ± 6.49 μm) were shorter than in the middle (363.76 ± 39.75 μm) and tip (390.69 ± 10.85 μm) regions. The exocuticle thickness showed minimal variation across segments with mean values of 6.06 ± 0.93 μm at the base, 3.83 ± 0.57 μm at the middle, and 3.56 ± 0.46 μm at the tip region (see Supplement).

**Figure 3.**
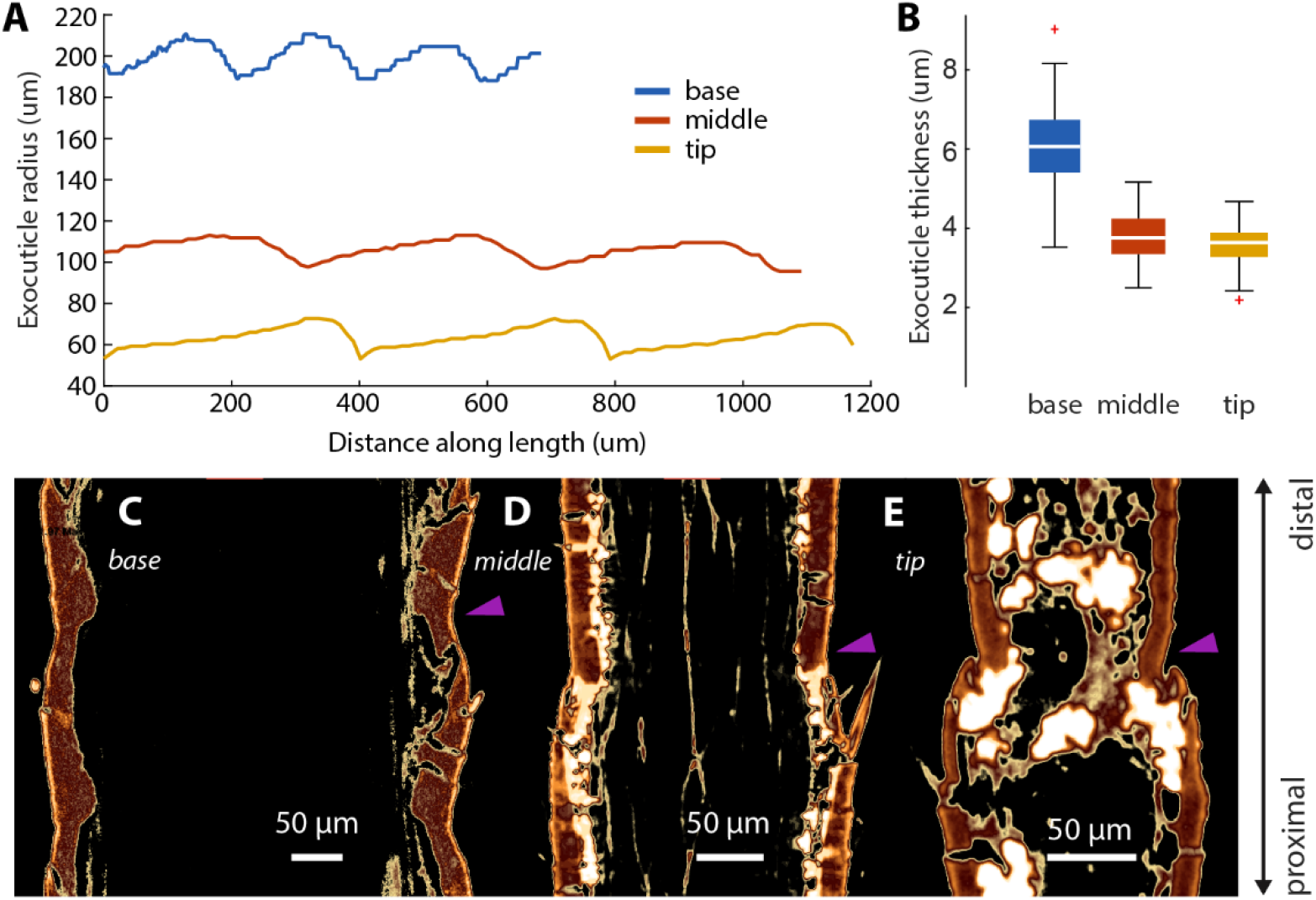
Morphology of the base, middle, and tip regions of the antenna reveals an invagination at the tip. (A) Radius variation within segments across the base (blue), middle (red) and tip (yellow). (B) Thickness for base, middle and tip regions. The intersection of (C) base, (D) middle, and (E) tip regions of antenna (purple arrows). The intersection between segments at the tip shows an invagination.

Micro-CT imaging revealed a unique invagination at the segment intersections in the tip region, distinguishing it from the base and middle regions. At these intersections, the cuticle surface folds inward and extends outward toward the next segment and therefore forms a S-shape geometry (**Figure 3E**). As this invagination is not found in the base and middle region, we hypothesized that it may serve a specialized function at the tip that facilitates large bending without buckling. Specifically, we speculate that during large bending due to a transverse load, the invagination may allow larger axial strain to develop at the segment intersection. Interestingly, we observed no specific discontinuity of the exocuticle at the intersection of the base and middle regions, however this must be interpreted with appropriate caution given the dehydration protocol for imaging (**Figure 3C, D**). Based on this observation, we hypothesized that the exocuticle may be continuous across segments and that the large deformation observed at the intersections is primarily caused by the decreasing radius, i.e. flexural stiffness, at these junctions.

### Finite element simulation supports a kinematic chain structure

To study the functional consequence of anatomical specializations in the antenna and test the hypothesis that the antenna functions as a continuous structure, we developed three region-specific finite element models of the base, middle, and the tip regions based on morphological data (**Figure 2;** see Materials and Methods). These models were used to simulate the bending response of the hypothesized continuous shell structure of the antenna. The finite element models were subjected to fixed-free beam boundary conditions. The nodes at the proximal end were fixed, while linear motion transverse to the neutral axis was applied to the distal end nodes (**Figure 4A**). The coordinates of the nodes in the ring at the intersections (red nodes in **Figure 4A**) were extracted, and the average coordinates of each ring were used to calculate joint angles between adjacent segments. Additionally, the nominal strain parallel to the antenna length in one element— positioned similarly to the marginal sensilla (Toh, 1977; Tominaga and Yokohari, 1982; Watanabe et al., 2018)—was recorded (blue dot in **Figure 4A**).

**Figure 4.**
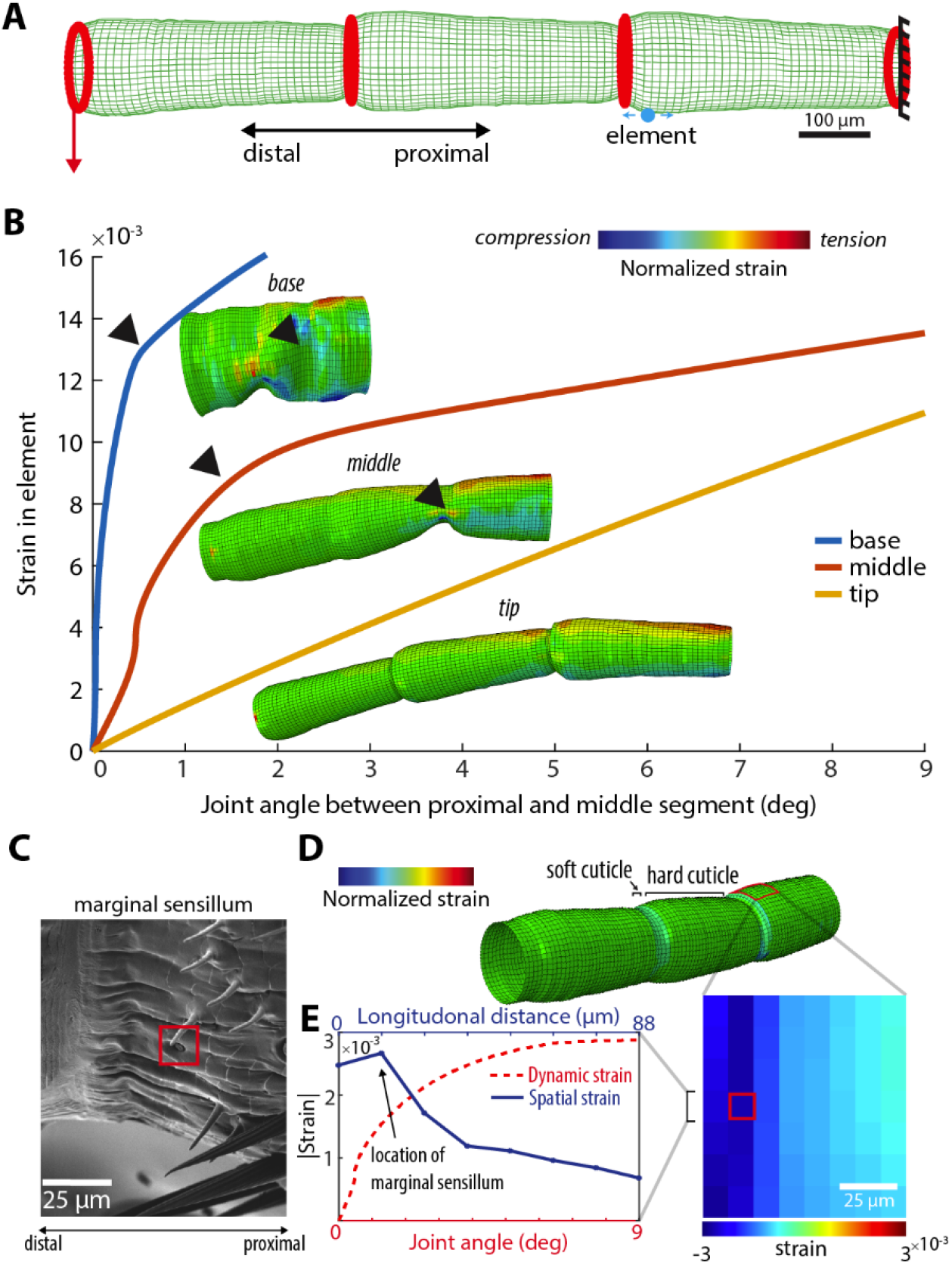
A finite element model of the antenna reveals biomechanical adaptions and estimates cuticular strain near marginal sensilla. (A) Simulation setup with fixed-free beam boundary conditions. The recorded element is marked by a blue dot, with nominal strain direction shown by the blue arrow. Red arrow: bending direction. Red rings: nodes used to calculate joint angles during bending. (B) Colormap: strain distribution, normalized individually based on the maximum and minimum strain values at the given time. Black arrow: buckling point. C) SEM image of marginal sensillum near the intersection between segments of the antenna base (10^th^ segment from scape). D) Hard cuticle strain field near marginal sensillum (red box) for intersegmental joint angle of 5° for the model of the middle of the antenna with soft cuticle between segments. E) Spatial and dynamic strain of the hard cuticle going through the approximate location of the marginal sensillum. The spatial strain is shown for a moderate intersegmental angle of 5 degrees.

As the joint angles between segments increased, buckling was observed in the base and middle regions, resulting in a nonlinearity in the strain-angle relationship (**Figure 4B**). In contrast, the tip region exhibited a linear relationship between element nominal strain and bending angle and showed no signs of buckling (**Figure 4B**). The observed buckling response in the base and middle regions aligns with the Brazier effect (Brazier, 1927), which describes the buckling of bent tubes (e.g., hollow plastic straws). During bending, longitudinal tension and compression tend to flatten or ovalize the cross-section of the tube. Once the curvature exceeds a critical value, instability occurs, leading to buckling (Calladine, 1983). In our simulations, buckling occurred at a critical segment angle of approximately 0.5° and 1.5° in the base and middle models, respectively.

However, in experiments, the observed segment angles were much larger: approximately 20° and 30° in the base and middle regions. This discrepancy indicates that under the continuous exocuticle assumption, the segment cannot achieve the large angles observed experimentally. Additionally, the folded movement between segments that was observed during antenna bending experiments was absent in the simulations. Therefore, our simulation suggests that the inclusion of soft cuticle intersections between segments is critical to model the flagellum response to large bending. These findings suggest that the large bending angles observed experimentally require an alternative explanation, such as material discontinuities in the membrane at the intersections.

### Invagination in tip region enhances flexibility

The FEM model of the tip exhibited greater segment bending angles compared to the base and middle models, in line with experimental data where the tip region withstood larger bending without structural failure (**Figure 4B**). Experiments and simulations suggest that the invagination in the tip region likely contributes to this enhanced flexibility for increased range of motion and improved conformity to mechanical stimuli. Taken together, our results suggest a novel morphological adaptation in the cockroach antenna.

### FEM estimates the strain field around marginal sensilla

Marginal sensilla are distributed along the flagellum and their location along the distal margin of segments suggests that they may function as proprioceptors that detect the articulation between segments (Schafer, 1971). They are identified by an elliptical depression in the cuticle, accompanied by a pea-shaped protrusion (**Figure 4C**). Marginal sensilla are categorized as a type of campaniform sensilla because they are thought to have similar functions (Schafer, 1971). To begin to describe the strain field around marginal sensilla and develop a neuromechanical model of the antenna, we repeated the FEM simulation but this time we explicitly modeled the “rigid-soft-rigid” architecture of the flagellum, as supported by our experimental data and kinematic chain model. To incorporate this modification, we reduced the elastic modulus of four layers of elements in the longitudinal direction (44 μm) at each segment intersection to 0.4 GPa. This lower elastic modulus was estimated based on tensile testing experiments on the flagellum of *P. americana* (Donley et al., 2022). Using the same boundary conditions described earlier and using the FEM model of the middle of the flagellum, we extracted the strain field near the marginal sensilla for a moderate intersegmental angle of 5° (**Figure 4D, Movie 4**). Our simulation revealed that marginal sensilla are located near the maximum strain of the hard cuticle strain gradient developed during transverse bending (**Figure 4E**). Further, our simulation showed how the strain near marginal sensilla scales with intersegmental angle (**Figure 4E**), thereby predicting the strain information available for antenna proprioception. The location of marginal sensilla could have important implications for understanding mechanisms of distributed strain sensing on insect antennae as the specialization of campaniform sensilla appears to derive primarily from their anatomical arrangement (Dickerson et al., 2020).

## DISCUSSION & CONCLUSION

Our study provides a critical step toward developing a detailed neuromechanical model of the antenna of a touch specialist, which could provide critical insights into mechanisms of touch sensation across insects. We showed that the filiform cockroach antenna is well approximated as a kinematic chain with localized bending at the segment intersection of soft cuticle. We also revealed distinct morphology across the base, middle and tip regions. Experiments and analytical modeling demonstrated that the maximum bending angle between successive segments increased from the base to the tip. Notably, while the middle region consistently fractured during cycling bending experiments, the tip region remained intact even under larger bending. Micro-CT imaging revealed an invagination at the segment intersections within the tip region, suggesting a morphological adaptation that enhances flexibility. Finite element analysis supported this notion, showing that the invagination enables larger bending without structural failure. The softer cuticle between segments appears to play a critical role in the antenna’s ability to undergo large deformation. By modeling these discontinuities in elastic modulus using FEM, we approximated the strain field across marginal sensilla. FEM showed how the strain near marginal sensilla scales with intersegmental angle, predicting the information available for antenna proprioception. Taken together, our findings provide insights into biological strategies for achieving both flexibility and structural integrity. Further, our findings could inspire the development of soft sensors by leveraging intricate geometries instead of softer materials for enhanced flexibility and resilience.

### Hybrid soft-rigid morphology of the antenna

Our simulations, based on the assumption of a continuous exocuticle, showed early buckling and failure in the base and middle regions, which was inconsistent with experimental observations. Similar breakage was observed when we reduced the elastic modulus of the exocuticle, suggesting that the intersectional membrane may be composed of a more elastic material compared to the more rigid exocuticle. This supports the idea of a contiguous ‘rigid-soft-rigid’ connection across segments, where soft cuticle at the intersections of rigid segments allows for intersegmental folding without structural failure. This hypothesis aligns with our observation of a transparent cuticle at the segment intersections. We propose that this transparent cuticle is composed of significantly lower modulus material, previously referred in literature as the soft or membranous cuticle (Andersen, 2010; Snodgrass, 1935). However, we cannot rule out that the apparent continuity of the exocuticle observed in micro-CT scans may be an artifact caused by dehydration of the sample. Cuticle mechanical properties are extremely sensitive to tanning (sclerotization) and water content: soft cuticle typically contains 40-75% water, whereas hardened cuticle (exocuticle) contains only ∼12% water (Vincent and Wegst, 2004). Dehydration before micro-CT scanning may have confounded the structural distinction between exocuticle and softer intersectional membrane, making the latter appear structurally continuous with the exocuticle. Supporting this, we observed that dried antennae from dead cockroaches and air-dried antennae became notably fragile (**Movie 5**). The fragility closely matched the mechanical failure seen in our continuous exocuticle simulations.

### Natural failure of the antenna

Cockroaches often experience partial antenna breakage due to aggressive behavior with conspecifics or through ontogeny. To further investigate this, we conducted the same bending experiment on a cockroach with a naturally broken antenna in the middle region. We observed that the fractured area had been sealed with hemolymph, forming a clot. During bending, movement in the damaged segment was significantly restricted, and when the bending force reached a critical threshold, a brittle fracture occurred (**Movie 6**). These observations suggest that when failure occurs in the intersectional membrane, hemolymph is released and solidifies into a clot, stabilizing the segment while maintaining overall structural continuity. Given that the cockroach antenna consists of approximately 150 segments, the immobilization of a single segment is unlikely to significantly impact its overall mechanical function.

### Application in soft robotics and neuromechanics

The findings of this study could provide inspiration for the development of soft tactile sensors. We found a unique geometry in the tip region of the cockroach antenna which enables large bending without buckling. This feature could be applied in soft robotic structures to enhance flexibility and durability. Unlike traditional kinematic chains in robotics, which may be limited to small-deformation scenarios, an antenna-inspired design could allow for more smooth, continuous motion while maintaining structural integrity (Demir et al., 2010). Additionally, our findings about the importance of water content in cuticle hint at potential of active modulation of mechanical properties for maintaining performance. Specifically, future soft robotic materials could benefit from tunable stiffness through controlled static pressure such as in hydrogels (Hedrick et al., 2024) or material phase transitions such as in shape memory alloys (Hedrick et al., 2024), polymers (Yuk et al., 2017) and electrohydraulic actuation (Kellaris et al., 2021). Such an approach could lead to the development of adaptive metamaterials (McClintock et al., 2021) and robotic structures that alter their mechanical properties in response to environmental conditions (Shah et al., 2021). Future research should explore how the segmental mechanics of insect antennae can be translated into functional prototypes for soft robotic systems to enable tactile perception in low-light environments. Finally, with advances in physics-based modeling (Todorov et al., 2012), our work could inspire more detailed neuromechanical models of the cockroach antenna with distributed sensing. By combining neural models of insect mechanoreceptors with a realistic physics simulation of the antenna, this work sets the stage to study how touch information across a population of distributed mechanoreceptors is represented at the neural level in the brain of an insect.

## Acknowledgements

We thank members of the Penn State Center for Quantitative Imaging for assistance with micro-CT imaging, and sample preparation, and image analysis. We thank Julie Anderson of the Microscopy Facility within the Huck Institute of the Life Sciences at Penn State for assistance with SEM imaging and image analysis.

## Author contributions

Conceptualization: L.M., P.M., K.J., J.M.M.; Methodology: L.M., J.M.M.; Formal analysis: L.M.; Investigation: L.M.; Writing: L.M., K.J., J.M.M; Supervision: K.J., J.M.M.; Funding acquisition: K.J., J.M.M.

## Data and resource availability

At the time of publication, all data and code will be deposited in Penn State ScholarSphere with a DOI.

## REFERENCES

Alba-Tercedor, J., Hunter, W. B. and Alba-Alejandre, I. (2021). Using micro-computed tomography to reveal the anatomy of adult Diaphorina citri Kuwayama (Insecta: Hemiptera, Liviidae) and how it pierces and feeds within a citrus leaf. Sci Rep 11, 1358.

Andersen, S. O. (2010). Insect cuticular sclerotization: a review. Insect Biochem Mol Biol 40, 166– 178.

Böröczky, K., Wada-Katsumata, A., Batchelor, D., Zhukovskaya, M. and Schal, C. (2013). Insects groom their antennae to enhance olfactory acuity. Proceedings of the National Academy of Sciences 110, 3615–3620.

Brazier, L. (1927). On the flexure of thin cylindrical shells and other “thin” sections. Proceedings of the Royal Society of London. Series A 116, 104–114.

Calladine, C. R. (1983). Theory of Shell Structures. Cambridge: Cambridge University Press.

Camhi, J. M. and Johnson, E. N. (1999). High-frequency steering maneuvers mediated by tactile cues: antennal wall-following in the cockroach. Journal of Experimental Biology 202, 631– 643.

Combes, S. A. and Daniel, T. L. (2003). Flexural stiffness in insect wings II. Spatial distribution and dynamic wing bending. Journal of Experimental Biology 206, 2989–2997.

Comer, C. M., Parks, L., Halvorsen, M. B. and Breese-Terteling, A. (2003). The antennal system and cockroach evasive behavior. II. Stimulus identification and localization are separable antennal functions. Journal of Comparative Physiology A 189, 97–103.

Cowan, N. J., Lee, J. and Full, R. J. (2006). Task-level control of rapid wall following in the American cockroach. Journal of Experimental Biology 209, 1617–1629.

Demir, A., Samson, E. and Cowan, N. J. (2010). A tunable physical model of arthropod antennae. IEEE International Conference on Robotics and Automation.

Dickerson, B. H., Fox, J. L. and Sponberg, S. (2020). Functional diversity from generic encoding in insect campaniform sensilla. Curr Opin Physiol 104743.

Donley, G., Sun, Y., Pass, G., Adler, P. H., Beard, C. E., Owens, J. and Kornev, K. G. (2022). Insect antennae: Coupling blood pressure with cuticle deformation to control movement. Acta Biomater 147, 102–119.

Hedrick, A., Kabutz, H., Smith, L., MacCurdy, R. and Jayaram, K. (2024). Femtosecond Laser Fabricated Nitinol Living Hinges for Millimeter-Sized Robots. IEEE Robot Autom Lett 9, 5449– 5455.

Kapitskii, S. V (1984). Morphology of the antenna of the male American cockroach Periplaneta americana. J Evol Biochem Physiol 20, 59–66.

Kellaris, N., Rothemund, P., Zeng, Y., Mitchell, S. K., Smith, G. M., Jayaram, K. and Keplinger, C. (2021). Spider-Inspired Electrohydraulic Actuators for Fast, Soft-Actuated Joints. Advanced Science 8,.

Lee, J., Sponberg, S., Loh, O. Y., Lamperski, A. G., Full, R. J. and Cowan, N. J. (2008). Templates and Anchors for Antenna-Based Wall Following in Cockroaches and Robots. IEEE Transactions on Robotics 24, 130–143.

Loudon, C., Bustamante, J. and Kellogg, D. W. (2014). Cricket antennae shorten when bending (Acheta domesticus L.). Front Physiol 5,.

McClintock, H. D., Doshi, N., Iniguez-Rabago, A., Weaver, J. C., Jafferis, N. T., Jayaram, K., Wood, R. J. and Overvelde, J. T. B. (2021). A Fabrication Strategy for Reconfigurable Millimeter-Scale Metamaterials. Adv Funct Mater 31,.

Mongeau, J.-M., Demir, A., Lee, J., Cowan, N. J. and Full, R. J. (2013). Locomotion-and mechanics-mediated tactile sensing: antenna reconfiguration simplifies control during high-speed navigation in cockroaches. Journal of Experimental Biology 216, 4530–4541.

Mongeau, J.-M., Demir, A., Dallmann, C. J., Jayaram, K., Cowan, N. J. and Full, R. J. (2014). Mechanical processing via passive dynamic properties of the cockroach antenna can facilitate control during rapid running. Journal of Experimental Biology.

Mongeau, J.-M., Sponberg, S. N., Miller, J. P. and Full, R. J. (2015). Sensory processing within cockroach antenna enables rapid implementation of feedback control for high-speed running maneuvers. Journal of Experimental Biology 218, 2344–54.

Okada, J. and Toh, Y. (2000). The role of antennal hair plates in object-guided tactile orientation of the cockroach (Periplaneta americana). Journal of Comparative Physiology A 186, 849– 857.

Okada, J. and Toh, Y. (2006). Active tactile sensing for localization of objects by the cockroach antenna. Journal of Comparative Physiology A 192, 715–726.

Pass, G. (2000). Accessory pulsatile organs: evolutionary innovations in insects. Annu Rev Entomol 45, 495–518.

Rajabi, H., Shafiei, A., Darvizeh, A., Gorb, S. N., Dürr, V. and Dirks, J.-H. (2018). Both stiff and compliant: morphological and biomechanical adaptations of stick insect antennae for tactile exploration. J R Soc Interface 15, 20180246.

Schafer, R. (1971). Antennal sense organs of the cockroach, Leucophaea maderae. J Morphol 134, 91–103.

Schafer, R. and Sanchez, T. V (1973). Antennal sensory system of the cockroach, Periplaneta americana: postembryonic development and morphology of the sense organs. Journal of Comparative Neurology 149, 335–353.

Schafer, R. and Sanchez, T. V (1976). The nature and development of sex attractant specificity in cockroaches of the genus Periplaneta. I. Sexual dimorphism in the distribution of antennal sense organs in five species. J Morphol 149, 139–157.

Schneider, D. (1964). Insect antennae. Annu Rev Entomol 9, 103–122.

Shah, D., Yang, B., Kriegman, S., Levin, M., Bongard, J. and Kramer-Bottiglio, R. (2021). Shape Changing Robots: Bioinspiration, Simulation, and Physical Realization. Advanced Materials 33,.

Snodgrass, R. (1935). Principles of Insect Morphology. McGraw-Hill.

Todorov, E., Erez, T. and Tassa, Y. (2012). MuJoCo: A physics engine for model-based control. In 2012 IEEE/RSJ International Conference on Intelligent Robots and Systems, pp. 5026–5033. IEEE.

Toh, Y. (1977). Fine structure of antennal sense organs of the male cockroach, Periplaneta americana. J Ultrastruct Res 60, 373–394.

Tominaga, Y. and Yokohari, F. (1982). External structure of the sensillum capitulum, a hygro-and thermoreceptive sensillum of the cockroach, Periplaneta americana. Cell Tissue Res 226, 309–318.

Vincent, J. F. V and Wegst, U. G. K. (2004). Design and mechanical properties of insect cuticle. Arthropod Struct Dev 33, 187–199.

Wainwright, S. A. (2020). Mechanical design in organisms. Princeton University Press.

Watanabe, H., Koike, Y., Tateishi, K., Domae, M., Nishino, H. and Yokohari, F. (2018). Two types of sensory proliferation patterns underlie the formation of spatially tuned olfactory receptive fields in the cockroach Periplaneta americana. Journal of Comparative Neurology 526, 2683– 2705.

Yuk, H., Lin, S., Ma, C., Takaffoli, M., Fang, N. X. and Zhao, X. (2017). Hydraulic hydrogel actuators and robots optically and sonically camouflaged in water. Nat Commun 8, 14230.

